# GeoFlow-V2: A Unified Atomic Diffusion Model for Protein Structure Prediction and De Novo Design

**DOI:** 10.1101/2025.05.06.652551

**Authors:** BioGeometry Team

## Abstract

We introduce GeoFlow-V2, a unified atomic diffusion model that seamlessly integrates structure prediction and de novo protein design across multiple biological modalities, including proteins, nucleic acids (DNA/RNA), and small molecules. The model’s core innovation lies in its unified architecture that natively handles both structure prediction (when provided with complete input sequences) and generative design (for partially or fully masked inputs) through a shared atomic diffusion process. Through the integration of structure conditioning constraints and the sequence design module, GeoFlow-V2 operates as a fully bidirectional framework, accepting both sequence and structural inputs while generating corresponding outputs. GeoFlow-V2 can also accommodate diverse experimental constraints and prior knowledge, which boosts performance and enables precise control over the folding and design process. Benchmarking against state-of-the-art methods demonstrates GeoFlow-V2’s strong performance in both structure prediction and de novo antibody design. We showcase GeoFlow-V2 ‘s versatility through protein design cases conditioned on diverse target modalities. To maximize accessibility, we engineer key functionalities of GeoFlow-V2 and provide a user-friendly web server at prot.design for non-commercial research use.

## 1. Introduction

Proteins serve as life’s essential molecular machinery, governing virtually all biological processes. The dual challenges of determining their three-dimensional structures (structure prediction) and engineering novel functional variants (protein design) represent foundational problems in computational biology. The field’s transformative potential was recently recognized by the 2024 Nobel Prize in Chemistry, awarded for groundbreaking advances in both protein structure prediction and computational protein design, highlighting the profound impact of these technologies on modern biological research.

Modern structure prediction methods, exemplified by AlphaFold 3 (Abramson et al., 2024), have achieved remarkable accuracy by leveraging deep learning approaches (LeCun et al., 2015) trained on extensive structural datasets. Concurrently, the field of protein design has entered a new era through the development of diffusion models (Bennett et al., 2024; Krishna et al., 2024; Watson et al., 2023) that can generate functional proteins with atomic-level precision, opening new possibilities for therapeutic development and synthetic biology.

Despite these parallel successes, current algorithms typically treat structure prediction and generative design as separate tasks. Recent efforts to repurpose structure prediction models for inverse design (Anishchenko et al., 2021; Cho et al., 2025; Krishna et al., 2024; Volk et al., 2023; Watson et al., 2023) have revealed untapped generative potential in these models, but remain limited by their original architectures.

In this technical report, we present GeoFlow-V2, a unified atomic diffusion model that bridges the gap between structure prediction and generative design across multiple biological modalities including proteins, nucleic acids, and small molecules. Our framework is built on the key insight that both structure prediction and design can be formulated as a conditional generation task where the model “inpaints” the input sequence and structure. E.g., in structure prediction, the model generates the full structure from a complete sequence; in protein binder design, the model generates the target-binder complex sequence and structure from the target sequence and structure. Under this formulation, we unify prediction and generation within a single, coherent diffusion generative framework through three key innovations: (1) pseudo protein sequences for unified protein generative modeling; (2) structure conditioning constraints and sequence co-design module, which enable the model to process both sequence and structural inputs and generate corresponding outputs; and (3) constraint-aware structure modeling that integrates experimental data and prior knowledge to guide the folding and design process.

The highlights of this report include:

- **A Unified Atomic Diffusion Framework**. GeoFlow-V2 establishes a unified architecture that bridges protein structure prediction and de novo design through a shared atomic diffusion process. The model handles both tasks natively–performing accurate structure prediction when given complete sequences while enabling generative design when provided with partially or fully masked inputs. This multimodal capability spans across key biological modalities including proteins, nucleic acids, and small molecules.
- **Versatile Constraint Support**. GeoFlow-V2 can incorporate diverse experimental constraints and prior knowledge to guide protein folding and design process.
- **Competitive Model Performance**. Comprehensive benchmarking demonstrates GeoFlow-V2’s strong performance across multiple tasks, including protein structure prediction and *de novo* antibody design. Our optimized lightweight variant, delivers 150-250x faster inference speeds than AlphaFold Multimer V2.3 for antibody structure predictions with even higher accuracy.
- **Accessibility**. A user-friendly web server is available at prot.design, providing three engineered functionalities of GeoFlow-V2, ranging from antibody-antigen structure prediction to *de novo* VHH design and protein binder design.

## 2 Overview

Our model architecture and training strategy primarily follow AlphaFold 3 (Abramson et al., 2024), with two distinct training configurations: for evaluation purposes, we use a training data cutoff of 2021-09-30 following Abramson et al. (2024); for production environments, we continuously train the full model with data through 2024-06-30. The key innovations of our method (Fig.1) are summarized in the following subsections.

**Figure 1.**
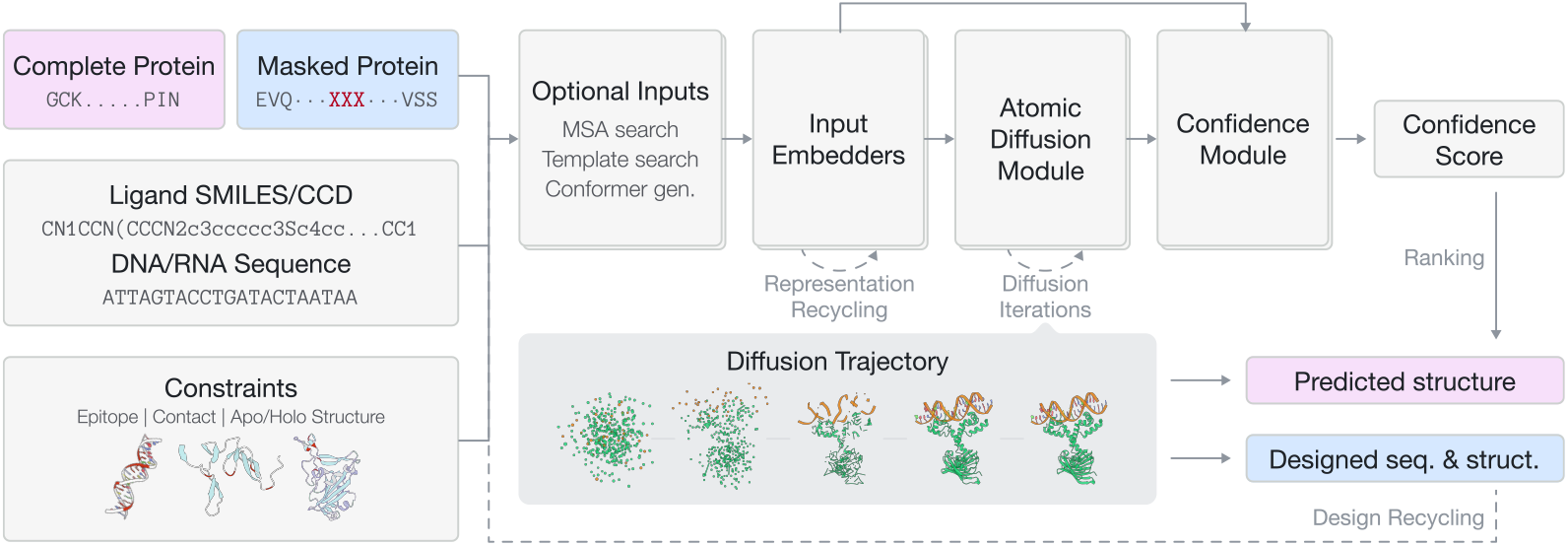
Overview of the GeoFlow-V2 model architecture and inference workflow. GeoFlow-V2 is a unified atomic diffusion model designed to seamlessly integrate both structure prediction and *de novo* protein design across multiple modalities, including DNA, RNA, molecules, and proteins. Key features include the ability to perform structure prediction when provided with complete protein sequences (and optionally sequences from other modalities), highlighted in purple, with the confidence module evaluating the accuracy of these predictions. For partially or fully masked proteins, GeoFlow-V2 performs *de novo* protein design, highlighted in blue, generating both structures and sequences using either its native co-design module or external inverse folding modules. The model accepts various constraint features, such as epitope constraints and target structure conditioning, enabling precise control over the folding and design process. Through the integration of the structure conditioning module and sequence design module, GeoFlow-V2 operates as a fully bidirectional framework, accepting both sequence and structural inputs while generating corresponding outputs. This capability enables design recycling for a second round of confidence score estimation, facilitating high-accuracy *in silico* screening of high-potential binders.

### 2.1 Unifying Structure Prediction and Generation

Recent advances in large-scale structure prediction models–including AlphaFold Multimer V2.3 (Evans et al., 2021), RoseTTAFold2 (Baek et al., 2023), AlphaFold 3 (Abramson et al., 2024), Protenix (Team et al., 2025), Chai-1 (team et al., 2024), and Boltz (Wohlwend et al., 2024)–have achieved remarkable success in biomolecular structure determination through training on extensive experimental and distilled structural data. While originally developed for structure prediction, recent studies have demonstrated these models’ latent capability for inverse structure generation, particularly in protein design (Anishchenko et al., 2021; Cho et al., 2025; Krishna et al., 2024; Volk et al., 2023; Watson et al., 2023). Crucially, we observe that structure prediction can be reformulated as a conditional generation task where the sequence specifies the desired output. We reason that by systematically altering the input conditions—*replacing complete sequences with (partially or fully) masked sequences*—we can unify structure prediction and generation within a single framework. The same architecture can thus seamlessly transition between the two capabilities through controlled conditioning. However, protein design additionally requires target structure conditioning and structure-conditioned sequence design, which the current architecture cannot natively support. We therefore introduce several key innovations to enable these essential features.

#### Pseudo Protein Sequence

The input protein sequence can exist in three states: (1) Complete sequence; (2) Partially masked sequence; or (3) Fully masked sequence. Masked residues are replaced with an “UNK” pseudo-residue containing only the four backbone atoms (C, N, O, and C*α*). The masking pattern is task-dependent (e.g., CDR residues for *de novo* antibody design) and are not required to be continuous. During training, we randomly apply masking using predefined schemes for either continuous sequence segments or spatially proximal residues, following AlphaFold 2’s cropping approach (Yang et al., 2023). The current implementation focuses solely on protein masking, and we leave the extension to other modalities for future work, but the model can condition on full sequences of other modalities (e.g., ligand).

#### Apo / Holo Structure Conditioning

Protein design applications—particularly protein binder design and motif scaffolding—frequently require target structure conditioning, especially for challenging targets (Bennett et al., 2023). To this end, we introduce a dedicated structure encoding module that encodes the coarse-grained distance map of the target structure. The structure is converted to a distance map in Angstrom, perturbed with Gaussian noise, and transformed into binned 2-dimensional one-hot features. This module operates orthogonally to the native template encoder, which handles searched and aligned structural templates. The integration of this module enables the model to accept raw structural inputs from either provided structures or its own outputs, while maintaining structural flexibility through Gaussian noise perturbation. This capability is particularly valuable in protein design workflows, where designed sequences / structures can be recycled for confidence verification and the iterative generation paradigm enables the refinement of previous designs.

#### Sequence Co-design Module

GeoFlow-V2 incorporates a novel sequence co-design module to generate amino acid types for masked residues in pseudo protein sequences. This module predicts sequence identities for masked regions at each diffusion step, conditioned on both structural representations from the diffusion module and amino acid predictions from the previous step. We note that off-the-shelf sequence design methods (Dauparas et al., 2022; Goverde et al., 2024) exist and can be readily integrated into our model. The integration of our structure conditioning module and sequence co-design module enables GeoFlow-V2 to operate as a fully bidirectional framework, accepting both sequence and structural inputs while generating both sequence and structural outputs.

### 2.2 Constraint Features

Prior structural knowledge—derived from wet-lab experiments (e.g., epitope mapping, alanine scanning mutagenesis data (Haynes et al., 2021; Moreira et al., 2007))—often provides critical constraints for protein structure prediction. For protein design tasks, we also frequently require both target structure conditioning and specified binding epitopes (hotspots). To incorporate these requirements, GeoFlow-V2 integrates three types of constraint features including epitope constraints, contact constraints, and Apo / Holo structure conditioning. Let *c*_*i*_ denote the *i*− th chain, *a*_*i, j*_ denote the *j* −th token in *i*− th chain, *d* denote a distance threshold, and 𝒰 denote the uniform distribution. The constraint features are defined as follows:

#### Epitope constraint

An epitope constraint is formally defined as the tuple (*a*_*i, j*_, *c*_*k*_, *d*), enforcing that the minimal distance of the token *a*_*i, j*_ to the chain *c*_*k*_ is smaller than *d* Angstrom, i.e.,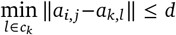.

Here, the distance between the tokens is defined as the Euclidean distance between representative atoms of corresponding tokens. During training, *d* is randomly sampled from 𝒰(6, 20).

#### Contact constraint

A contact constraint is formally defined as the tuple (*a*_*i, j*_, *a*_*k,l*_, *d*), enforcing that the distance between the token *a*_*i, j*_ and the token *a*_*k,l*_ is smaller than *d* Angstrom, i.e., ∥*a*_*i, j*_ − *a*_*k,l*_ ∥ ≤ *d*. During training, *d* is randomly sampled from 𝒰 (6, 30).

#### Structure conditioning constraint

As discussed in Section.2.1, this constraint is the noise-perturbed distance map extracted from target structures, which are then binned into one-hot features.

We encode epitope constraints and contact constraints using Gaussian smearing (Schütt et al., 2017):

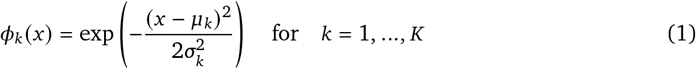

where *u*_*k*_ is calculated as linearly spaced centers and *σ*_*k*_ as the uniform bandwidth:

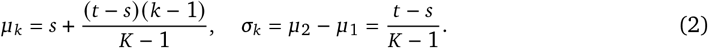

The hyper-parameters are set to (*s* = 6, *t* = 20, *k* = 6) for epitope constraints and (*s* = 6, *t* = 30, *k* = 6) for contact features. The mask value is set to −100. These continuous representations provide gradient sensitivity to distance thresholds and outperform the one-hot encoding. For structure conditioning constraints, noise-perturbed distance maps are binned into 0−4, 4−8, 8 − 16, *>* 16, with an additional bin for mask values.

To prevent over-reliance on constraints, each constraint type is independently activated with 20% probability during training (replaced with pre-defined mask values when inactive), supplemented with constraint dropout. Epitope and contact constraints are stochastically generated from ground-truth complexes, where the number of constraints per sample follows a geometric distribution (*p* = 1/3).

For structure conditioning, we randomly partition ground-truth complexes into binder and target groups–systematically assigning antigens as targets in antibody-antigen systems–and only target group structures are encoded as constraints.

## 3. Protein Structure Prediction

We evaluate GeoFlow-V2’s structure prediction performance across three distinct tasks: (1) antibody-antigen complex prediction using a curated low-homology dataset, (2) protein-ligand docking on the PoseBusters benchmark (Buttenschoen et al., 2024), and (3) antibody structure prediction on an independently curated test set. While the model demonstrates robust general protein modeling capabilities, we particularly emphasize antibody-related assessments–highlighting GeoFlow-V2’s specialized focus as a unified framework for protein design, especially for *de novo* antibody design. These targeted validations verify the structural accuracy required for reliable generative applications.

### 3.1 Antibody-Antigen Structure Prediction

#### Setup

We assess prediction performance on a rigorously curated low-homology test set following AlphaFold 3’s interface validation protocol (Supplementary Section 5.8). The dataset was constructed by collecting all PDB antibody-antigen complexes released between 2024-06-30 and 2025-01-30, cropping antibodies to their Fv regions, and filtering complexes to contain 256-1024 residues for computational efficiency. This process yielded 104 high-quality antibody-antigen complexes for benchmarking. We compare GeoFlow-V2 against several state-of-the-art approaches: the AlphaFold 3 replicas **Protenix** (Team et al., 2025), **Chai-1** (team et al., 2024), and **Boltz** (Wohlwend et al., 2024). The direct AlphaFold 3 comparison was precluded by license restrictions. We also include **AlphaFold Multimer V2.3** and **GeoFlow-V1** (BioGeom, 2024) as baselines. For diffusion-based methods, we generate 50 predictions per target (10 random seeds × 5 diffusion samples each), while AlphaFold Multimer V2.3 predictions are obtained by ensembling 5 model weights with 10 random seeds each. All predictions are evaluated using the DockQ score (Basu and Wallner, 2016) under three scenarios: Top-1 (single highest-ranked prediction), Top-10 (best prediction among the top 10 ranked outputs), and Oracle (best possible prediction across all samples). We define the DockQ success rate as the percentage of antibody-antigen complexes with DockQ *>* 0.23.

#### GeoFlow-V2 achieves state-of-the-art performance on antibody-antigen docking

Fig.2A (left panel) illustrates the comparative docking performance across evaluated methods. GeoFlow-V2 demonstrates superior accuracy, achieving a 45.19% top-1 success rate across all three evaluation scenarios–surpassing both AlphaFold 3-derived replicas and established baselines. Key observations and discussions are as follows.

- GeoFlow-V2 establishes new state-of-the-art performance, while Protenix, Chai-1, and GeoFlowV1 form a secondary tier.
- Despite these advances, antibody-antigen docking remains fundamentally challenging, with substantial space for improvement even for top-performing methods.
- We hypothesize that further gains could be achieved through: (1) Inference-time scaling (Ma et al., 2025) (additional random seeds / MSA searches); (2) Enhanced noise sampling strategies (Yang et al., 2025); and (3) More accurate confidence estimation modules.

#### Boosting Docking Performance with Constraints Guiding

We next evaluate how different constraint features enhance GeoFlow-V2’s docking accuracy on low-homology antibody-antigen sets using three DockQ metrics: acceptable (DockQ *>* 0.23), medium (DockQ *>* 0.49), and high (DockQ *>* 0.80).

Six evaluation settings are compared: (1) Blind docking using only sequence inputs, (2) One Contact constraint with a single antibody-antigen residue pair within either 15Å or 25Å C*α* distance, (3) One Epitope constraint with a random antigen residue within 8Å of the antibody, (4) Four Epitopes with four such antigen residues, and (5) Holo Antigen structure conditioning constraint, while still requiring full complex prediction from random initialization. Key observations (Fig.2B) and discussions are as follows:

- Structural constraints substantially improve docking performance, with four epitope constraints boosting the top-1 acceptable success rate from 45% to 75%, demonstrating particular value when prior knowledge from experimental data or design hotspots are available. An example is visualized in Fig.2C.
- Holo antigen constraints show unique advantages for high-accuracy predictions (DockQ *>* 0.80), despite potential information leakage from the bound structure, making them especially useful for antibody optimization (Cai et al., 2024) and antibody design (Bennett et al., 2024) scenarios where bound structures might be available.
- Four-epitope constraints achieve the best overall performance in acceptable and medium accuracy ranges, confirming the model’s ability to precisely follow specified binding hotspots–a critical capability for *de novo* protein design applications. These results collectively establish the GeoFlow-V2 as an effective and flexible tool for both constrained and unconstrained antibody-antigen docking tasks.

### 2.3 Protein-Ligand Structure Prediction

Following the Chai-1 (team et al., 2024) evaluation protocol, we define prediction success as achieving a ligand root mean square deviation (RMSD) below 2Å from the ground truth structure. Our benchmarking compares GeoFlow-V2 against three established baselines: Chai-1, AlphaFold 3, and RoseTTAFold AllAtom (Krishna et al., 2024), using published results from their respective papers. For GeoFlow-V2 evaluation, we analyze the top-ranked prediction across five model seeds, with each seed generating five diffusion samples. As shown in Fig. 2A (right panel), GeoFlow-V2 achieves a 77% success rate on the protein-ligand structure prediction task. This result demonstrates the ability of the model to model protein-ligand interactions, which is crucial for designing ligand-binding proteins where precise molecular recognition is essential.

**Figure 2.**
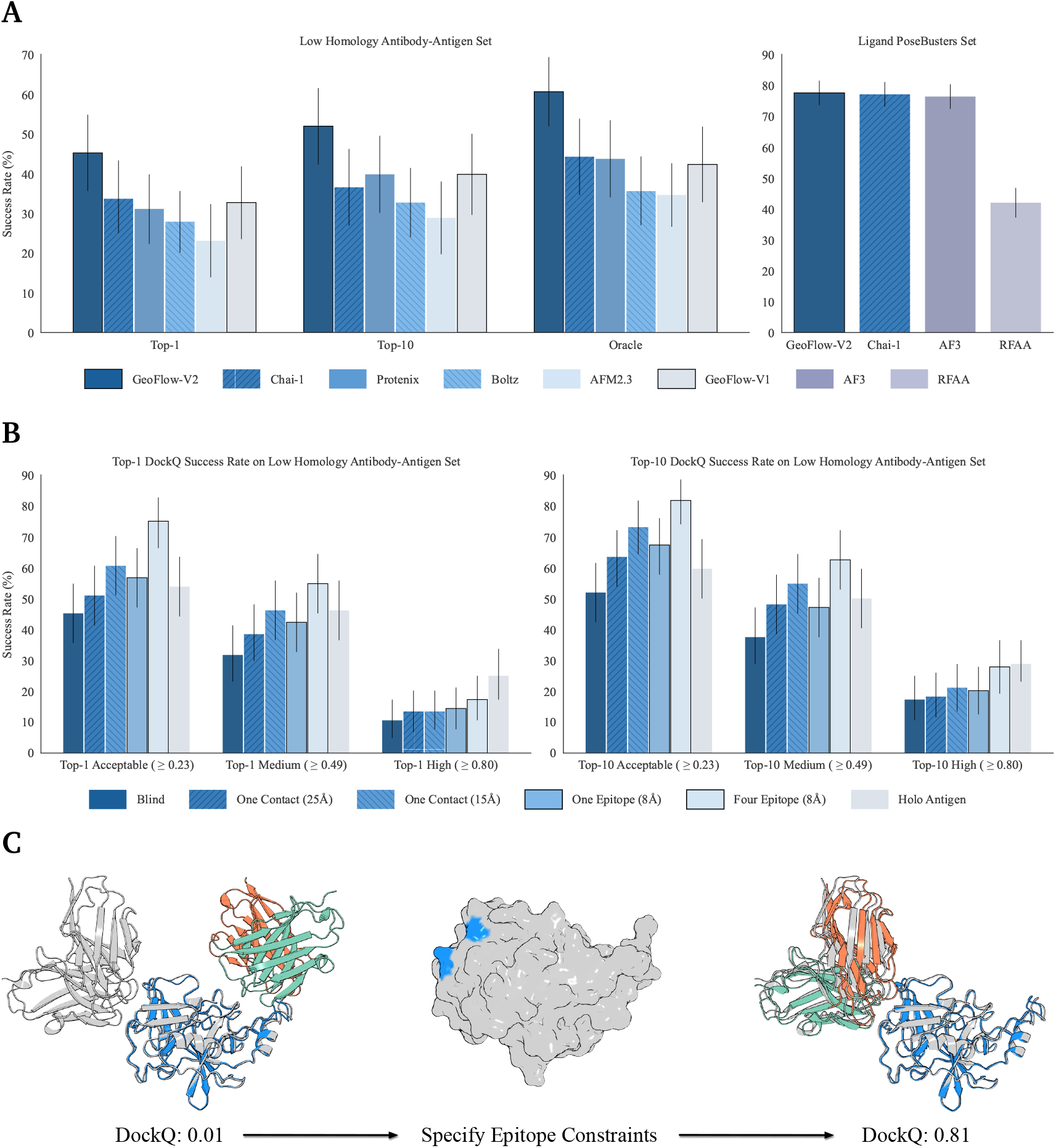
Performance comparison across several benchmarks. For all panels, the bar heights represent the mean values across test cases, with error bars indicating 95% confidence intervals obtained through 10,000 bootstrap resamples for the antibody-antigen set and exact binomial distribution for PoseBusters set. **(A)** Comparison of protein complex structure prediction performance across different models. **Left panel**: Percentage of antibody-antigen complexes with DockQ *>* 0.23 on a low-homology antibody-antigen test set, evaluated under three prediction scenarios: Top-1 (top-ranked prediction), Top-10 (best prediction among top-10 ranked predictions), and Oracle (best possible predictions). **Right panel**: Performance on the Ligand PoseBusters benchmark set. Success rates represent the percentage of samples with pocket-aligned ligand RMSD < 2 Å. The results of AF3 were calculated using provided predictions and the results of RFAA were taken from AF3’s paper. The scores are calculated on the top-ranked prediction based on its confidence scores. **(B)** Benchmark performance of GeoFlow-V2 on a low-homology antibody-antigen test set, with varying levels of docking constraint features. Bars are grouped by different docking accuracy levels (Acceptable, Medium, High) measured on top-ranked predictions (left panel) and Top-10-ranked predictions (right panel). **B***)Blind*: Vanilla GeoFlow-V2 without constraints, taking only antibody and antigen sequences as inputs. *One Contact (15Å / 25Å)*: We randomly select one antibody-antigen residue pair within 15Å / 25Å C*α* distance and provide this pair along with the distance threshold as template features to the model. *One Epitope (8Å) / Four Epitopes (8Å)*: We define the epitope as one (or four) random antigen residue(s) with C*α* atom within 8Å of the antibody chain, and then provide these residues as template features to the model. *Holo Antigen*: We provide the bound structure of the antigen chains as template features to the model. Note the model is still required to generate the whole antibody-antigen structures from random Gaussian initialization. **(C)** A cherry-picked example of antibody-antigen structure prediction for PDB ID 8VGI with and without epitope constraints. Coloring scheme: Gray for ground-truth antibody-antigen complexes; Blue for predicted antigens and epitope residues; Green and Orange for predicted heavy and light chains.

### 3.3 Ultra-fast Antibody Structure Prediction

Accurate antibody structure prediction remains challenging, especially for CDR loop modeling (Ruffolo et al., 2023). Current models (Abanades et al., 2022; Ruffolo et al., 2022) typically rely on limited specialized datasets like SAbDab (Dunbar et al., 2014), which contains approximately 10,000 antibody structures. Inspired by AlphaFold 2 monomer distillation techniques (Jumper et al., 2021), we hypothesize that training a lightweight GeoFlow-V2 variant on the OAS database’s (Olsen et al., 2022) millions of paired antibody sequences could simultaneously improve both prediction accuracy and computational efficiency, with details provided next.

#### Lightweight GeoFlow-V2 Training

We develop a lightweight variant of GeoFlow-V2 specialized for antibody structure prediction named as GeoFlow-V2-ab, by reducing the number of input embedding layers from 48 to 8 while maintaining the core architecture. The model follows a two-phase training protocol as visualized in Fig.3D. First, pretraining is performed exclusively on the SAbDab dataset with a 2024-06-30 cutoff date. This dataset is rigorously curated by removing structures with incomplete Fv regions, excluding entries with resolutions worse than 4Å, cropping to Fab regions, and clustering antibodies at a minimum CDR similarity threshold of 0.8.

**Figure 3.**
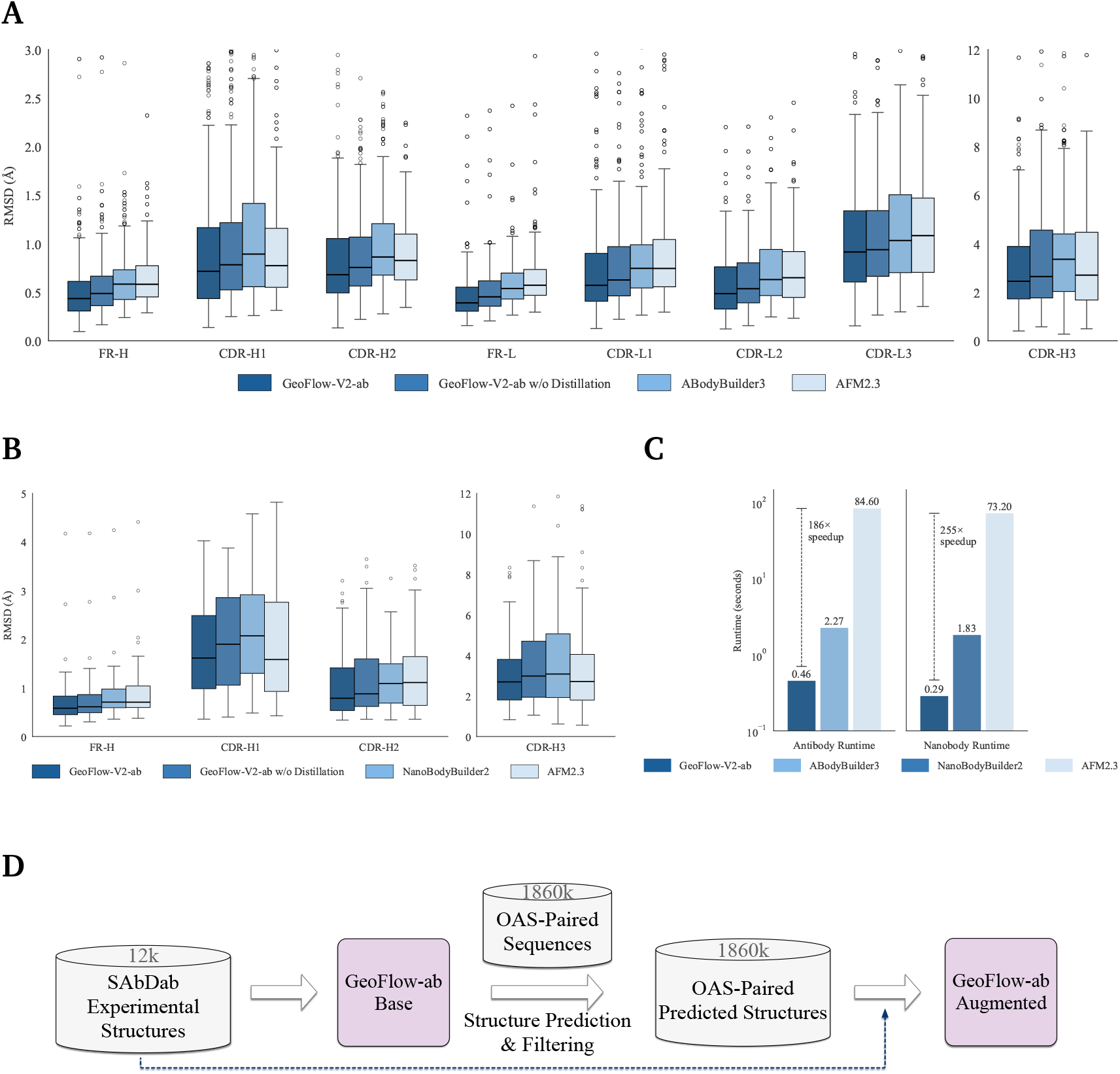
Performance and runtime comparison for antibody structure prediction. The root-mean-square deviation (RMSD) values were calculated over backbone heavy atoms after aligning the corresponding framework residues. The box plots display the median (center line), interquartile range (IQR, box boundaries), with whiskers extending to 1.5 × IQR. Outliers beyond 1.5 x IQR are shown as individual points. **(A)** Benchmark performance of GeoFlow-V2-ab, GeoFlow-V2-ab (without OAS-paired distillation), ABodyBuilder3, and AlphaFold-Multimer V2.3 (AFM2.3) on antibody structure prediction (n = 205 test cases). Due to the high structural variability and typically larger deviations in CDR-H3, we present it separately (right panel) to allow for a more detailed comparison across methods, while the left panel shows RMSD distributions for framework regions and other CDRs (H1, H2, L1–L3). **(B)** Benchmark performance of GeoFlow-V2-ab, GeoFlow-V2-ab (without OAS-paired distillation), NanoBodyBuilder2, and AFM2.3 on nanobody structure prediction (n = 80 test cases). **(C)** Antibody (left panel) and nanobody (right panel) prediction runtimes (log scale) for GeoFlow-V2-ab, ABodyBuilder3 / NanoBodyBuilder2, and AFM2.3. Bar heights show average runtime for predicting one structure in seconds, with dashed lines indicating speedup factors relative to AFM2.3. GeoFlow-V2-ab demonstrates consistent 150-250× faster runtime compared to AFM2.3 while maintaining competitive or higher accuracy. **(D)** Workflow for developing specialized GeoFlow-V2 for antibody structure prediction. The model is first pre-trained on SAbDab dataset, which is used to create the OAS-paired distilled dataset. The lightweight GeoFlow-V2 is then re-trained from scratch using a balanced 1:1 mixture of the original SAbDab data and this new distilled set.

The trained model predicts structures for approximately 1.86 million paired antibody sequences from the OAS database. We filter predictions using a confidence score threshold of 0.70, retaining high-quality outputs to create the OAS-paired distilled dataset. The lightweight GeoFlow-V2 is then re-trained from scratch using a balanced 1:1 mixture of the original SAbDab data and this new distilled set.

#### Performance Evaluation

For rigorous evaluation, we construct a low-homology test set containing 285 antibodies–205 conventional antibodies and 80 nanobodies–released after the training cutoff (post 2024-06-30) from clusters absent in the training data. We compare GeoFlow-V2-ab against ABodyBuilder3 (Kenlay et al., 2024) and AlphaFold Multimer V2.3 for antibody structure prediction, and include NanoBodyBuilder2 (Abanades et al., 2023) and AlphaFold Multimer V2.3 as baselines for nanobody structure prediction. The comparison against IgFold (Ruffolo et al., 2023) was precluded by license restrictions. We evaluate all methods by calculating backbone heavy-atom (N, C*α*, C, O) RMSD values after framework residue alignment, with separate analyses for antibodies (Fig.3A) and nanobodies (Fig.3B). Key observations include:

- All methods demonstrate high accuracy in predicting framework structures for both heavy and light chains, achieving median sub-angstrom RMSD values for CDR1 and CDR2 regions. However, this precision does not extend equally to CDRH1 and CDRH2 in nanobodies, likely due to limited nanobody training data availability.
- GeoFlow-V2-ab Distilled establishes new state-of-the-art performance. While the median improvement over the naive model is modest, the distilled version significantly reduces outlier predictions evidenced by lower 1.5×IQR ranges, demonstrating more robust performance.
- we observe that AlphaFold Multimer V2.3, despite its computational intensity and MSA requirements, is frequently outperformed by specialized baselines in multiple evaluation settings. This suggests dedicated antibody structure predictors may be preferable for routine applications where runtime efficiency is crucial.

#### Runtime Analysis

Fig.3C compares prediction runtimes (log scale) for antibody (left panel) and nanobody (right panel) structure prediction across three methods: GeoFlow-V2-ab, ABodyBuilder3 / NanoBodyBuilder2, and AlphaFold Multimer V2.3. All benchmarks were conducted on identical hardware (single NVIDIA A100 GPU). GeoFlow-V2-ab achieves consistent 150–250× faster inference than AlphaFold Multimer V2.3 while maintaining competitive or better accuracy. The combination of computational efficiency and maintained accuracy suggests GeoFlow-V2-ab is particularly suited for scenarios requiring rapid structure generation, such as high-throughput antibody screening. It can also aid antibody optimization workflows when structures are undetermined (e.g., structure-based developability prediction).

## 4. De Novo Antibody Design

To assess the performance of GeoFlow-V2 in *de novo* antibody design, we focus on *in silico* evaluations across three critical dimensions: (1) Structure Generation Performance–quantifying the model’s ability to generate structurally valid antibodies with optimal paratope geometry and favorable interaction patterns; (2) Binder Discrimination Performance–benchmarking the model’s capability to differentiate true binders from non-binders based on wet-lab data; (3) Virtual Screening Success Rates–assessing the model’s ability to produce promising candidates that meet *in silico*-defined binding criteria. Based on these computational insights, *de novo* VHH and antibody libraries have been designed against several therapeutic-relevant targets and subsequently synthesized as oligo pools for downstream validation. Experimental results will be reported in a future update to this technical report.

### 4.1. Structure Generation Performance

#### Setup

This task evaluates the model’s capability to generate structurally valid antibodies by measuring (1) whether the generated antibody structures adhere to the specified hotspots; and (2) whether the generated antibody structures adopt CDR-loop-mediated interactions. To this end, we curate a benchmark set named GeoFlow benchmark, consisting of 7 published therapeutic targets (Shanehsazzadeh et al., 2023a) (IL17A, ACVR2B, FXI, TSLP, IL36R, TNFRSF9, C5) for antibody design and 3 in-house targets for nanobody design. We note that the 7 published antibody targets have corresponding reference antibody-antigen complex structures as well as 1,243 binding data points obtained via inverse folding and SPR (Shanehsazzadeh et al., 2023a), while the 3 nanobody targets do not have any reference nanobody binding poses.

We benchmark GeoFlow-V2 against RFAntibody (Bennett et al., 2024), the current state-of-the-art antibody design method with full design workflow and experimental validation, which is built upon the RFdiffusion framework (Watson et al., 2023). While other machine learning-based antibody design approaches exist, direct comparison was infeasible due to several limitations: some methods (Luo et al., 2022) require known binding poses as input, while others (Wang et al., 2024) impose non-commercial licensing restrictions.

Unless specified otherwise, we follow Bennett et al. (2024) and adopt the standardized humanized VHH framework (h-NbBcII10_FGLA_) (Vincke et al., 2009) as the template for nanobody (VHH) design. For conventional antibody design, we utilize the Trastuzumab framework (Cho et al., 2003). This approach aligns with real-world design campaigns, where predefined frameworks are typically employed rather than native frameworks from reference antibodies. Importantly, it avoids potential model memorization of native reference antibody frameworks while providing a robust starting point for candidate developability by leveraging well-characterized frameworks. For each target antigen, we (1) select five binding-critical hotspot residues which are used by both methods; (2) generate 1,000 de novo designed structures using both GeoFlow-V2 and RFAntibody (following the instructions from its official github repository); and (3) the CDR lengths are set to {L1:8-13,L2:7,L3:9-11,H1:7,H2:6,H3:8-13} for conventional antibodies and {H1:7,H2:6,H3:8-16} for nanobodies under the Chothia definition.

#### Trastuzumab Antibody Sequences

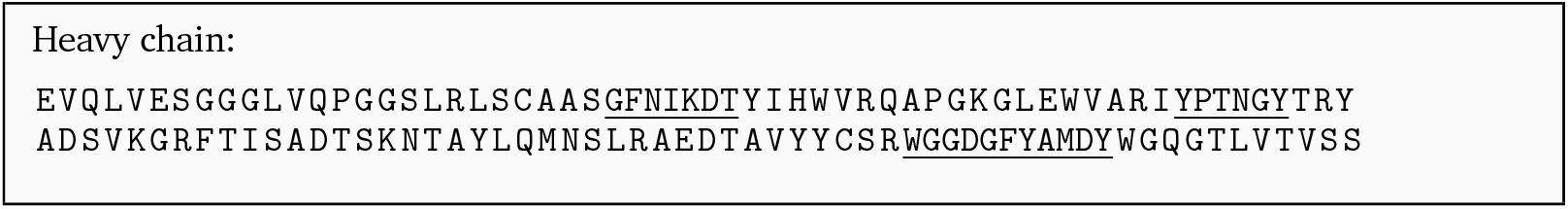
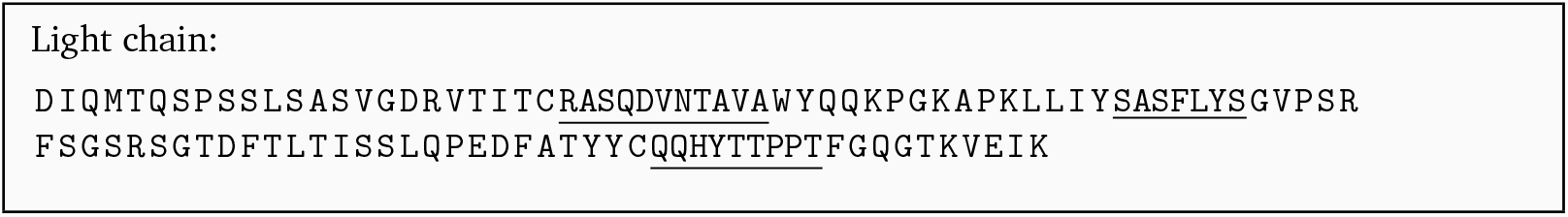

#### h-NbBcII10_FGLA_ VHH Sequence

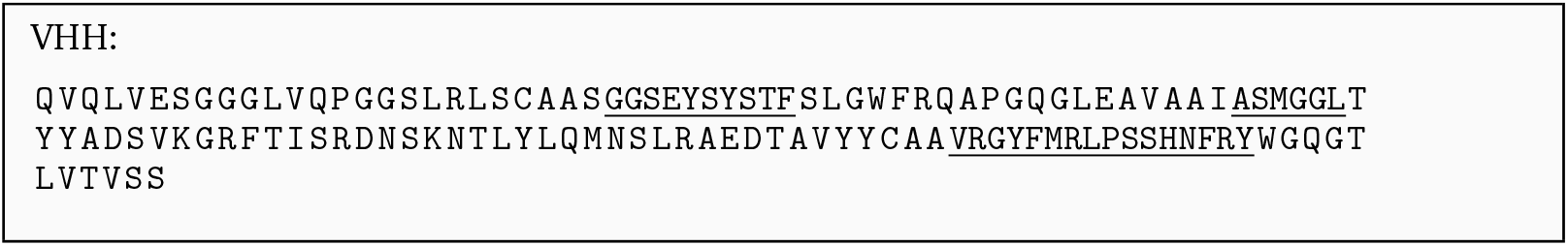

We evaluate structure generation performance using five metrics:

- **Hotspot** Pass Rate: The percentage of designs where ≥3 specified hotspot residues (60% of total) maintain minimal C*α* distances *<* 9 Å to the antigen. This relaxed criterion balances stringency with practical design feasibility.
- **CDR Interaction** Pass Rate: The percentage of designs with CDR-mediated antigen interfaces. While framework-mediated binding (Fig.4A) occurs naturally (Zavrtanik et al., 2018) and is also reported in Bennett et al. (2024), we exclude such cases as they usually require framework redesign for high affinity (Yamamoto et al., 2025), which is beyond our scope. We use the following relaxed criteria to determine if the structure adopts the CDR-mediated interactions. For each CDR region, we define the CDR interaction criterion by a tuple *name, n, d*, where *name* denotes the name of the CDR region (e.g., HCDR3), *n* denotes the minimal interacting residues for a CDR to be considered as interacting, and *d* denotes the interacting distance threshold. A residue is considered as interacting if it has minimal C*α* distance *< d* Å to the antigen.
- The relaxed interaction criterion for antibody is:

**Figure 4.**
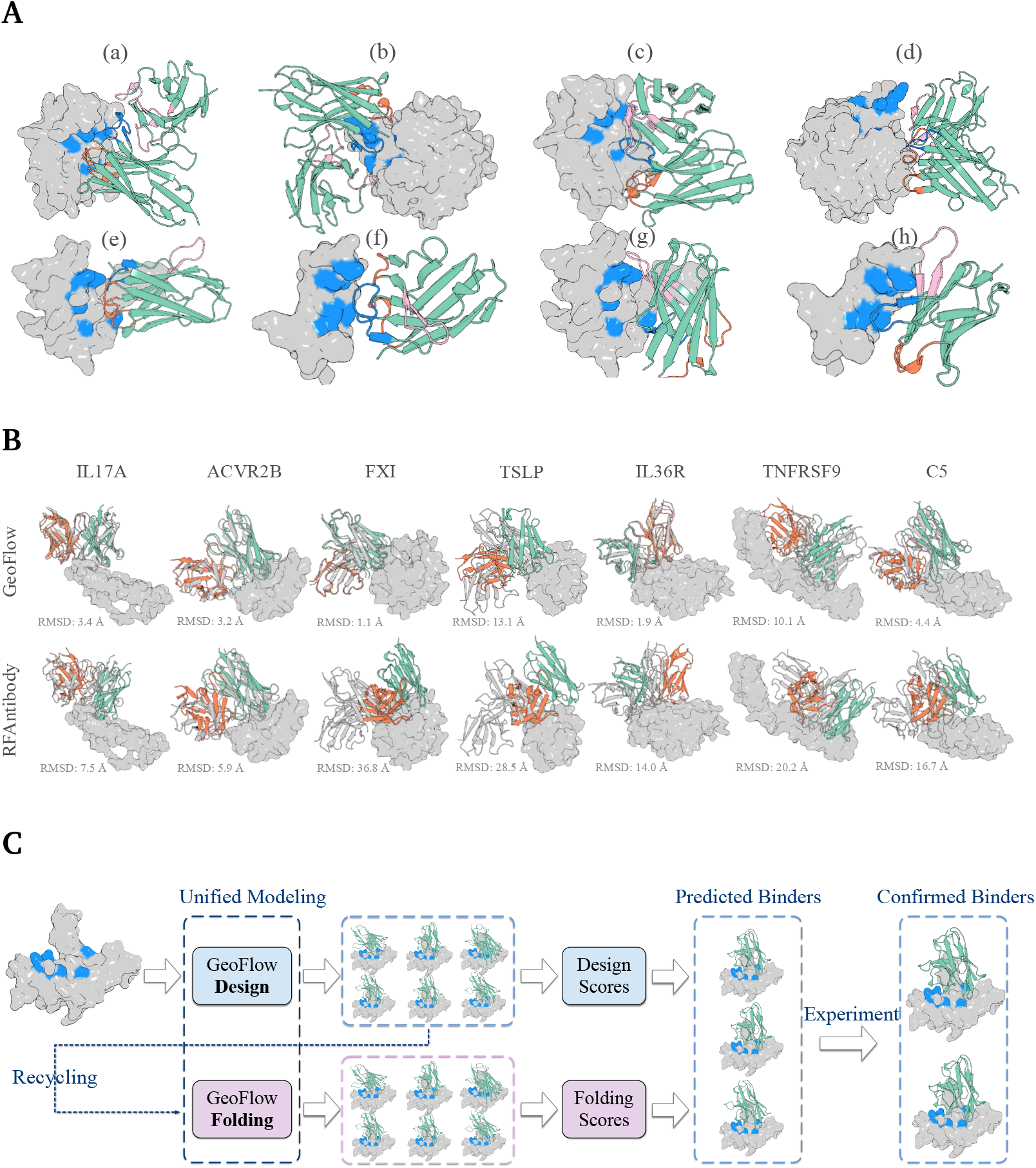
Overview of Antibody Design and Validation Workflow. (**A**) Representative structures from structure generation evaluation. Antigen representation: Gray surface with blue highlights for specified hotspot residues. Coloring scheme: Blue for CDR-H3; Orange for CDR-H1/H2; Pink for CDR-L (antibodies) or framework region 2 (nanobodies). (a & b) Antibodies with successful hotspots targeting. (c & d) Antibodies with failed hotspots targeting. (e & f) Nanobodies with CDR-mediated interactions. (g & h) Nanobodies with frameworkmediated interactions. **(B)** Designed antibody structures with minimal C*α* RMSD to reference poses across seven targets, generated using the Trastuzumab template framework. Coloring scheme: Gray for native antibody-antigen complexes; Green for designed heavy chains; Orange for designed light chains. **(C)** Workflow of design system. Starting from the target structure (hotspots highlighted in blue), the GeoFlow-V2 generates de novo designed structures and sequences of antibody candidates, which are recycled into GeoFlow-V2 for refolding (with complete antibody sequences). Confidence scores from both design outputs and folding outputs are used to predict whether a design will bind. Finally, the predicted binders are validated through experiments to identify confirmed binders.

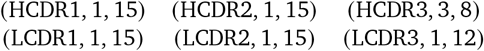

The relaxed interaction criterion for nanobody is:

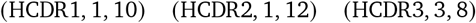

- **Overall** Pass Rate: The percentage of designs meeting both hotspot and CDR-interface criteria while being sterically clash-free (per AF3 standards (Abramson et al., 2024)).
- **Framework Recovery Rate**: The percentage of designed structures (averaged among targets) achieving framework C*α* root-mean-square deviation (RMSD) below specified thresholds. The RMSD values are calculated over shared framework residues (Chothia numbering) after antigen superposition, which measures the orientation fidelity of designed structures to reference binding poses. Note: Higher RMSD values do not necessarily indicate worse designs, as valid alternative binding modes may exist. This serves as an efficient computational proxy for structural validation before experimental characterization.
- **Diversity**: The percentage of designed structures exhibiting novel binding modes. Generated structures are clustered using a threshold of 5Å framework C*α* RMSD, and Diversity is calculated as the ratio of the number of clusters to the total number of designs. We note that this metric is correlated to the number of generated samples. For the sake of computational efficiency and to avoid the sample size effect, we randomly subsample 100 structures from 1,000 generated structures and report the mean statistic over multiple rounds.

#### Interaction Pattern Evaluation

We compare GeoFlow-V2 and RFAntibody performance on the GeoFlow benchmark using three metrics: **Hotspot, CDR Interaction**, and **Overall**. As shown in Fig. 5A, key findings include:

**Figure 5.**
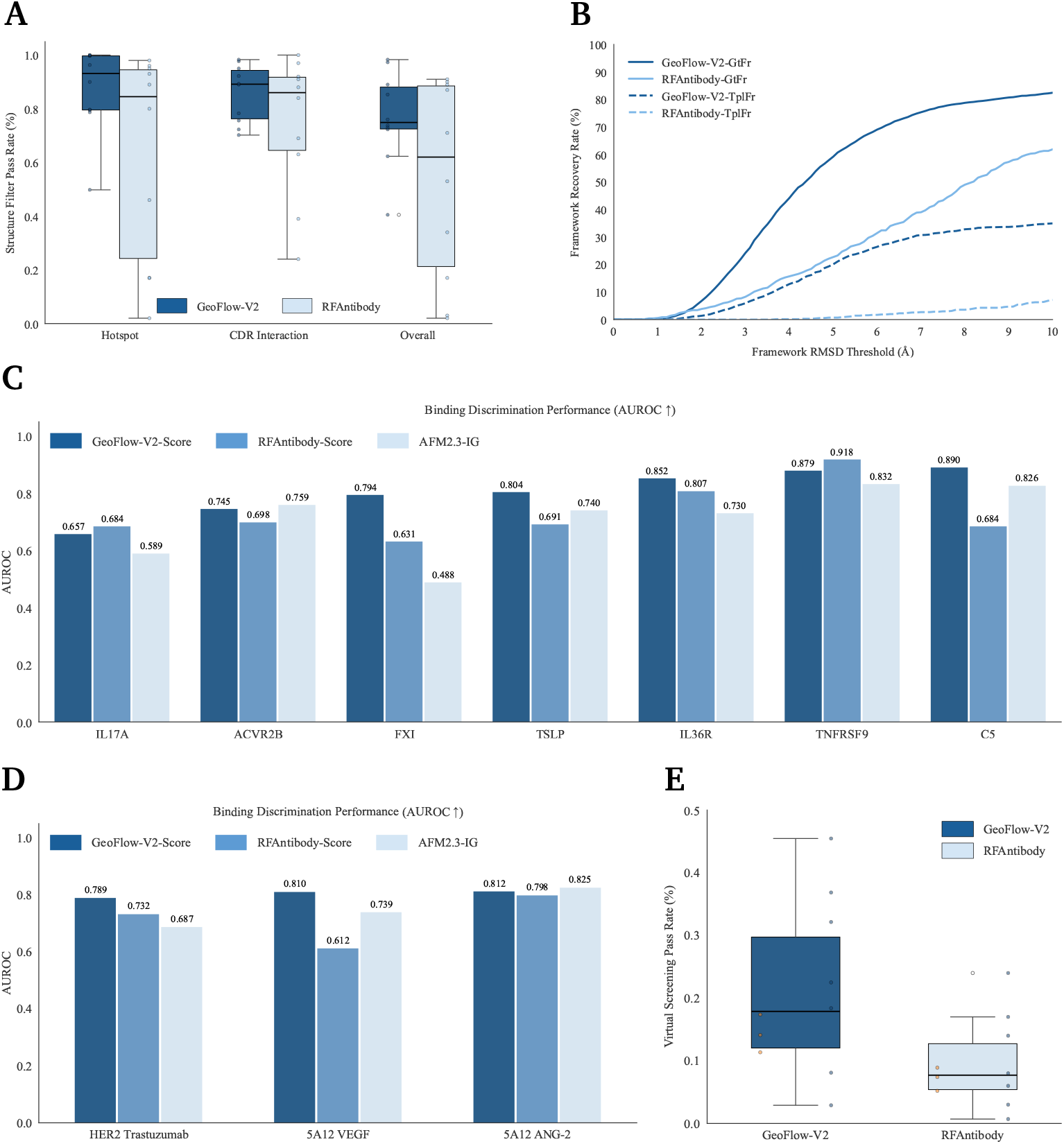
*In-silico* performance of GeoFlow-V2 and baselines across multiple benchmarks. **(A)** Performance of GeoFlow-V2 and RFAntibody on curated structure quality benchmark set, consisting of 7 published targets (IL17A, ACVR2B, FXI, TSLP, IL36R, TNFRSF9, C5) for antibody design and 3 in-house targets for nanobody design. Box plots show the distribution of averaged structure filter pass rates for each target, with whiskers extending to 1.5 x IQR. Outliers beyond 1.5 x IQR are shown as individual points. Individual data points (averaged pass rate for each target) are overlaid as dodged dots. Metrics are as follows. **Hotspot**: percentage of designs that adhere to the specified hotspots. **CDR Interaction**: percentage of designs with *CDR-loop-mediated* interfaces. **Overall**: percentage of designs that pass both Hotspot and CDR Interaction filters. The detailed definitions of these criteria can be found in the main text. **(B)** Performance of GeoFlow-V2 and RFAntibody on recovering reference antibody’s framework orientation under various RMSD thresholds. The RMSD values were calculated over shared antibody framework residues between designed antibodies and reference antibodies under the Chothia numbering scheme after aligning the target (antigen) residues. Solid curves (GtFr) show recovery results for designs that use the ground-truth antibody frameworks, while the dashed curves (TplFr) show results for designs that use a shared template antibody (or nanobody) framework. **(C & D)** Performance of GeoFlow-V2, RFAntibody, and AlphaFold-Multimer V2.3 initial guess (AFM2.3-IG) on the binder / non-binder discrimination task across 10 targets. The bar heights represent the AUROC values. **(E)** In silico virtual screening success rates with GeoFlow-V2-Scorer of GeoFlow-V2 and RFAntibody on the curated target set (7 published targets and 3 in-house targets). Box plots show the distribution of success rates for each target, with whiskers extending to 1.5 x IQR. The individual data points are visualized as horizontally-dodged dots, with nanobody targets highlighted in orange and antibody targets shown in blue.

- Both methods achieve high median pass rates (*>* 0.8) across ten targets, demonstrating strong baseline performance.
- However, GeoFlow-V2 shows superior robustness, particularly on three challenging targets–one in-house nanobody and two antibody targets (ACVR2B and FXI)–where RFAntibody underperforms. This may reflect RFAntibody’s reported sensitivity to precise hotspot selection.
- While both methods maintain median overall pass rates above 0.5, GeoFlow-V2 delivers more robust and consistent results across all targets.

#### Framework Recovery Evaluation

We assess GeoFlow-V2 and RFAntibody on 7 antibody targets from IgDesign (Shanehsazzadeh et al., 2023a), excluding nanobody targets due to unavailable reference poses. Two framework scenarios are evaluated: (1) *GtFr*: Using reference antibody frameworks (direct RMSD calculation via target alignment); (2) *TplFr*: Using template frameworks (Trastuzumab; RMSD calculated heuristically over Chothia-numbered shared residues). As shown in Fig. 5B, key findings include:

- Both methods achieve strong recovery with reference frameworks (*GtFr*), even at stringent thresholds.
- GeoFlow-V2 significantly outperforms RFAntibody with template frameworks (*TplFr*), demonstrating its superior generalizability for real-world antibody design without reference antibody frameworks.

We present representative structures that fail to adhere to specified hotspots or adopt framework-mediated interfaces in Fig.4A, and present the best-performing (lowest C*α* RMSD to reference poses) designed antibody structures across seven antibody targets in Fig. 4B, generated using the Trastuzumab template framework. These results demonstrate GeoFlow-V2’s capability to produce high-quality antibody candidates, enabling efficient in silico screening and subsequent experimental validation.

#### Diversity Evaluation

Both GeoFlow-V2 and RFAntibody demonstrate the ability to generate diverse binding modes across all ten targets, as measured by novel structural clustering (Tab. 1). These results were obtained using fixed hotspot configurations and default diffusion hyperparameters. In practical applications, diversity could be further enhanced through the strategic selection of multiple hotspot configurations and optimization of diffusion hyper-parameters. While RFAntibody shows better diversity metrics, we attribute this difference to the integration of structure prediction tasks in GeoFlow-V2, which may impose beneficial structural inductive bias (as indicated by better structure generation performance) that slightly reduce conformational exploration–a trade-off that can be modulated by adjusting the task ratio during training.

**Table 1.** The percentage of designed structures exhibiting unique binding modes across ten targets.

|  | Nb-1 | Nb-2 | Nb-3 | TSLP | FXI | IL36R | TNFRSF9 | C5 | ACVR2B | IL17A |
| --- | --- | --- | --- | --- | --- | --- | --- | --- | --- | --- |
| GeoFlow-V2 | 7% | 16% | 14% | 7% | 17% | 12% | 29% | 26% | 11% | 31% |
| RFAntibody | 8% | 19% | 19% | 26% | 20% | 19% | 40% | 57% | 21% | 51% |

### 4.2 Binder Discrimination Performance

#### Setup

While high-throughput in silico antibody generation can produce millions of candidates, wetlab validation remains bottlenecked by experimental throughput. To bridge this gap, computational pre-screening is essential for prioritizing high-probability binders before costly wet-lab testing. In this section, we evaluate GeoFlow-V2’s ability to discriminate true binders from non-binders using ten published benchmark datasets with wet-lab validated binding data. The statistics of ten datasets are summarized in Tab.2.

**Table 2.**
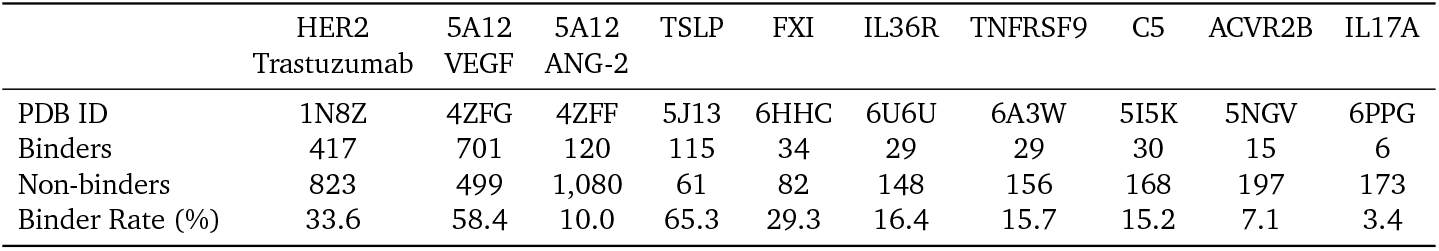
Summary of binder/non-binder datasets used for discrimination evaluation.

|  | HER2<br>Trastuzumab | 5A12<br>VEGF | 5A12<br>ANG-2 | TSLP | FXI | IL36R | TNFRSF9 | C5 | ACVR2B | IL17A |
| --- | --- | --- | --- | --- | --- | --- | --- | --- | --- | --- |
| PDB ID | 1N8Z | 4ZFG | 4ZFF | 5J13 | 6HHC | 6U6U | 6A3W | 5I5K | 5NGV | 6PPG |
| Binders | 417 | 701 | 120 | 115 | 34 | 29 | 29 | 30 | 15 | 6 |
| Non-binders | 823 | 499 | 1,080 | 61 | 82 | 148 | 156 | 168 | 197 | 173 |
| Binder Rate (%) | 33.6 | 58.4 | 10.0 | 65.3 | 29.3 | 16.4 | 15.7 | 15.2 | 7.1 | 3.4 |

- **HER2 Trastuzumab** (Shanehsazzadeh et al., 2023b): This benchmark comprises 1,240 designs (417 SPR-confirmed binders vs 823 non-binders) generated through zero-shot AI-based CDR engineering of trastuzumab targeting HER2 (PDB ID: 1N8Z).
- **5A12 VEGF & 5A12 ANG-2** (Minot and Reddy, 2024). These two datasets are derived from combinatorial mutagenesis of CDR-H2 (9 residues) and CDR-L1 (8 residues) in the 5A12 antibody, validated via yeast display with FACS sorting. The VEGF library contains 642,080 variants (375,267 binders, 58.4% rate), with our benchmark subset comprising 701 binders and 499 non-binders reflecting the original distribution. The ANG-2 library includes 711,912 variants (13,227 binders, 1.86% rate), subsampled to 120 binders and 1,080 non-binders for balanced evaluation. (PDB ID: 4ZFG & 4ZFF)
- **IgDesign Datasets** (Shanehsazzadeh et al., 2023a): This benchmark comprises seven therapeutic targets from the IgDesign study, featuring SPR-validated binding data for designs generated via two strategies: (1) HCDR-3-only redesign and (2) full HCDR-123 redesign within native antibody frameworks. The curated dataset contains 1,243 designs (258 binders and 985 non-binders) with various target-specific binder rates.

We benchmark GeoFlow-V2 with the following baselines: **RFAntibody-Score**, the filtering module finetuned from RoseTTAFold2 in the state-of-the-art antibody design method RFAntibody (Bennett et al., 2024); and **AlphaFold Multimer V2.3-initial guess**, a variant of AlphaFold Multimer V2.3 (Evans et al., 2021) equipped with the *initial guess* mechanism similar to Bennett et al. (2023). Note that unlike structure prediction, binder discrimination for antibody design remains understudied in computational biology, with no established benchmarks or standardized metrics. Our selection represents the most relevant approaches available. The prediction protocol is as follows:

- **GeoFlow-V2-Score** evaluates designed antibody-antigen complexes by taking both the antibody and antigen sequences and five binding-critical hotspot residues as input epitope constraints. The model performs full structure prediction of the designed complexes and generates multiple confidence scores similar to AlphaFold 3 (Abramson et al., 2024), including a self-consistency metric that quantifies the agreement between the designed structures and GeoFlow-V2’s predictions. While both design and folding stages produce confidence scores, this evaluation exclusively uses the folding-stage scores. We visualize this procedure in Fig.4C.
- **RFAntibody-Score**. We utilize the official implementation of RFAntibody. The model was fine-tuned from RoseTTAFold 2, which can predicts the structures of designed antibody-antigen complexes and produce a set of confidence scores, such as interface PAE (Predicted Aligned Error), target-aligned antibody RMSD, and framework-aligned CDR RMSD.
- **AlphaFold Multimer V2.3-Initial Guess** provides a MSA-free assessment of designed antibody-antigen complexes by leveraging AlphaFold’s structure prediction capability. The implementation takes the designed complexes as templates (chain level) and initialization of the structure module,

while strategically masking all CDR regions to prevent information leakage from the designed structures. The model generates predicted complex structures and multiple confidence scores, such as interface PAE, target-aligned antibody RMSD, binder-aligned RMSD, etc. We implement this model using OpenFold’s (Ahdritz et al., 2024) codebase.

#### Evaluation Results

We choose interface PAE scores as our primary binding metrics for all models, based on empirical performance optimization. We treat reference antibody-antigen structures as “designed” structures. While we recognize that such practice may not fully reflect real-world design scenarios, this approach remains justified for our benchmark context as the datasets primarily consist of designs generated through either (1) inverse folding or (2) combinatorial mutations starting from the reference structures. We exclude any designs with length discrepancies from their reference structures. Performance is quantified using the Area Under the Receiver Operating Curve (AUROC) metric, with results presented in Fig.5C and Fig.5D. Key findings include:

- All methods show predictive power, but none dominates across all targets. GeoFlow-V2 achieves the most consistent performance, while AlphaFold Multimer V2.3 shows the highest variance.
- GeoFlow-V2 outperforms in robustness, maintaining stable accuracy across diverse targets.
- Manual inspection reveals that GeoFlow-V2 more accurately recovers designed binding poses and maintains hotspot contacts compared to RFAntibody and AlphaFold Multimer V2.3. This may explain GeoFlow-V2’s superior robustness.

Binder discrimination performance represents a critical bottleneck for achieving high success rates in *de novo* antibody design. While further wet-lab validation data will be essential for developing more accurate models, our evaluation demonstrates that GeoFlow-V2-Score serves as a promising computational screening tool for antibody design pipelines.

### 4.3 Virtual Screening Success Rates

We evaluate the end-to-end performance of GeoFlow-V2 and RFAntibody in designing high-quality antibody candidates by measuring their virtual screening pass rates – the percentage of designs predicted to bind via our GeoFlow-V2-Score metric. The assessment uses the same ten targets from our structure generation benchmarks, with consistent design protocols as described in Section 4.1. For each target, we select five binding-critical hotspot residues and generate 1,000 *de novo* designed antibody structures and sequences. For RFAntibody, we augment each structure with 10 ProteinMPNN (Dauparas et al., 2022)-designed sequences and keep the top-scoring sequence. All designs are evaluated identically using GeoFlow-V2-Score’s interface PAE metric (Section 4.2), with the 8.5 Å threshold empirically determined to separate binders from non-binders. The virtual screening pass rate (visualized in Fig.5E) quantifies each method’s ability to produce viable candidates that meet this binding criterion. Key findings include:

- GeoFlow-V2 achieves higher median success rates (0.179) compared to RFAntibody (0.077), demonstrating superior robustness across all ten targets. While this comparison uses GeoFlowV2 ‘s scoring module and might be biased, the consistent performance gap suggests meaningful improvements in design quality.
- We observe substantial variation in success rates across targets, with both methods struggling on challenging cases like TNFRSF9 (GeoFlow-V2: 0.029, RFAntibody: 0.007). These results suggest the need for target-specific threshold adjustments and higher experimental throughput in real-world scenarios.
- The systematically lower pass rates for nanobody targets highlight limitations in current training data, emphasizing the requirement for dedicated nanobody optimization in future algorithmic developments.

These *in silico* evaluations demonstrate GeoFlow-V2 ‘s capability in generating structurally valid antibodies with optimized paratope geometry, discriminating high-affinity binders, and enriching promising candidates through virtual screening.

## 5. Unified Generative Modeling for Protein Binder Design

### 5.1 Overview

GeoFlow-V2 provides a unified framework for protein binder design through atomic-level modeling of molecular interactions. The approach supports diverse target modalities–including proteins, nucleic acids (DNA/RNA), and small molecules–while accommodating flexible design constraints such as (1) specified binding sites or interaction modes through epitope and contact constraints; and (2) partial structural knowledge of either target or binder via structure conditioning constraint features. The following sections discuss potential use cases across different target classes.

### 5.2 Design of Flavin-binding Proteins

Flavin cofactors (Mewies et al., 1998) such as FAD and FMN are essential redox-active groups in natural enzymes. Designing proteins capable of binding and potentially utilizing flavins could pave the way for novel enzymes with redox activity, or for applications in biocatalysis (Tong et al., 2023), biosensing (Bitzenhofer et al., 2022), and synthetic metabolic pathways (Duan et al., 2025). GeoFlowV2 can generate diverse, high-quality potential binders by directly specifying FAD or FMN as a design condition. It also provides the capability to redesign known motifs by specifying regions for modification, which can be beneficial in contexts where a balance between functional retention and structural variation is desired. Below we present an example of an FAD binder candidate generated by GeoFlow-V2. The resulting protein–ligand complex exhibits a well-packed and structurally reasonable binding conformation, with numerous intermolecular interactions.

**Figure 6.**
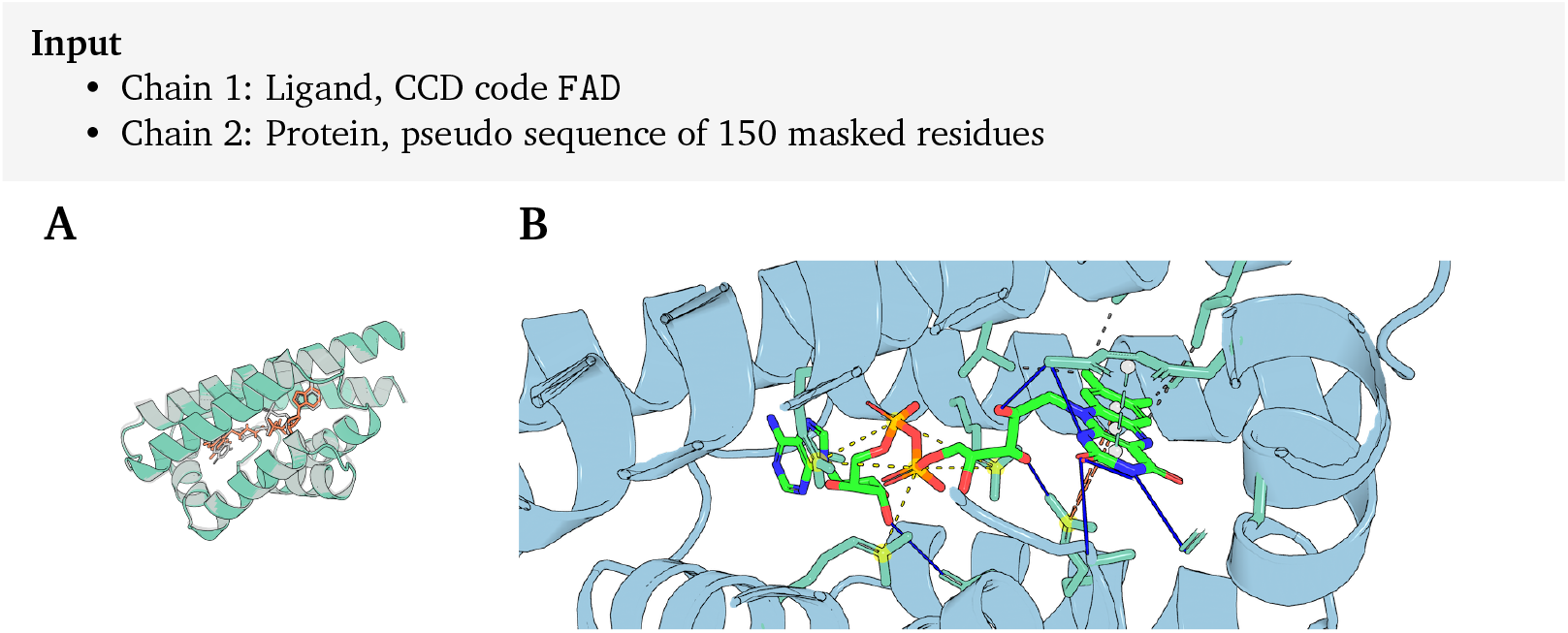
Example of a flavin-binding protein designed by GeoFlow-V2. **(A)** Illustration of an FAD binder candidate designed by GeoFlow-V2. The design exhibits a self-consistency RMSD of 0.689 and a consistency confidence score of 1.58. The designed binder structure is shown in green, the designed FAD binding conformation in orange, and the re-predicted binder structure and binding conformation in gray. **(B)** Zoomed-in view of the interactions between the designed FAD binder and FAD, generated using PLIP (Adasme et al., 2021). Hydrophobic interactions, hydrogen bonds, *π*–stacking (parallel), *π*–cation interactions, and salt bridges are represented by black dotted, blue solid, green dotted, orange dotted, and yellow dotted lines, respectively.

### 5.3 Design of OKT3-Masking Peptide

OKT3 is a murine monoclonal antibody of the immunoglobulin IgG2a isotype (Norman, 1995), targeting CD3, a signal complex that initiates T cell activation and determines the specificity of the immune response (Menon et al., 2023). Designing a short peptide that binds to OKT3, i.e., a masking peptide, may offer a potential means of modulating immune responses. GeoFlow-V2 can effectively leverage known antibody structures (Kjer-Nielsen et al., 2004) to design diverse binding peptides in key loop regions. Below we present examples of OKT3-masking peptide generated by GeoFlow-V2.

**Figure 7.**
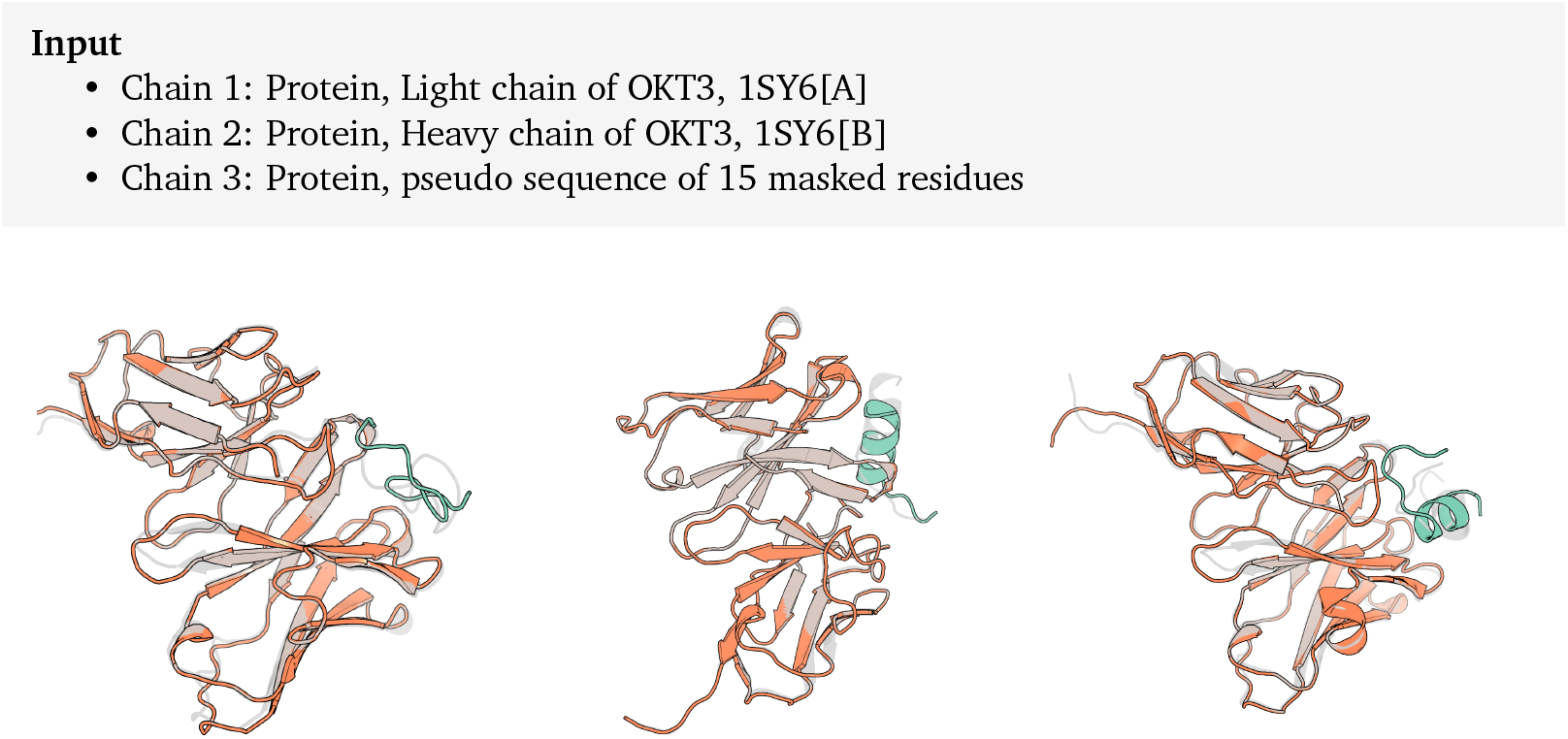
Examples of OKT3-masking peptide designed by GeoFlow-V2. The designed peptide structure is shown in green, the designed OKT3 conformation in orange, and the re-predicted complex structure and binding conformation in gray. Although structural information of the target is provided, GeoFlow-V2 does not rigidly fix the target conformation; instead, it allows for a certain degree of structural flexibility to facilitate the formation of a more plausible binding mode.

### 5.4 Design of Competitive Binder for NhaR-mediated Transcriptional Activation

NhaR is a transcriptional activator protein in Escherichia coli (strain K12) that promotes the expression of the nhaA Na+/H+ antiporter gene by binding to a specific DNA sequence upstream of the gene, thereby enabling the cell to respond to changes in pH and salt concentration (Rahav-Manor et al., 1992). Designing a synthetic protein that competes with NhaR for binding to this regulatory DNA region could block NhaR-mediated transcriptional activation, offering a potential strategy for modulating associated physiological processes. GeoFlow-V2 enables *de novo* design of DNA-binding proteins directly from target sequence information. GeoFlow-V2 allows for the specification of a target hotspot position as a design constraint. While the model generally adheres to the constraint with a high likelihood, alternative binding mode may still be explored when structurally favorable. Below we present examples of competitive binder generated by GeoFlow-V2.

**Figure 8.**
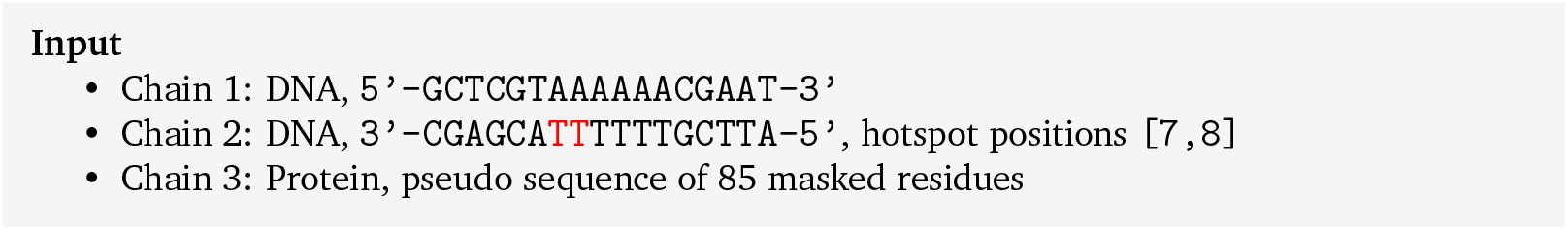
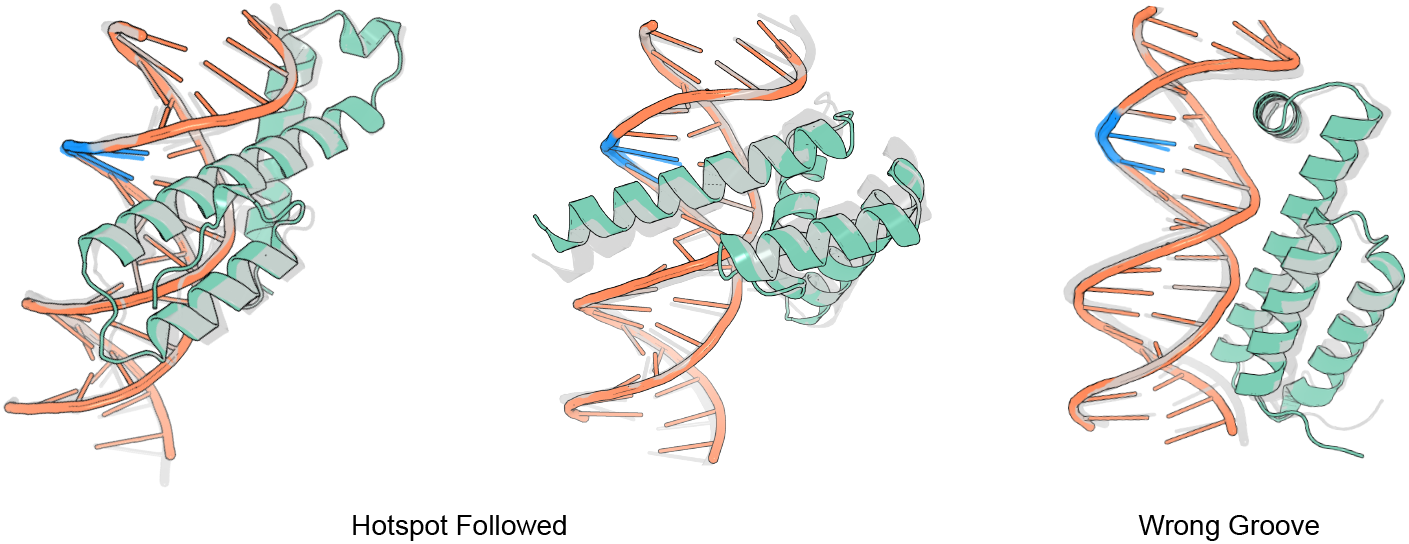
Examples of competitive binders for nhaR-mediated transcriptional activation, designed by GeoFlow-V2. Coloring scheme: Green for the designed peptide structures; Blue for hotspot regions; Orange for the designed DNA conformations; Gray for the re-predicted complex structures and binding conformations. Left: examples of binders designed by GeoFlow-V2 targeting the groove specified by the input hotspot. Right: an example where the binder docks to an alternative groove. While GeoFlow-V2 generally adheres to the specified hotspot with high probability, alternative binding modes may still occur.

## 6. Conclusion, Limitations and Future Directions

We have presented GeoFlow-V2, a unified atomic diffusion framework that establishes a novel paradigm for bridging structure prediction and generative design. While the model demonstrates strong performance and provides a unified generative framework, we also acknowledge that GeoFlow-V2 has some limitations. First, the architecture requires pre-defined atom configurations for generative design (e.g., C, N, O, C*α* for masked protein residues), and cannot yet handle small molecule generation and nucleic acid generation. Second, as a pure all-atom model without the frame-based inductive biases used in AlphaFold 2, it occasionally produces ligand conformations with incorrect chirality. Third, there remains substantial space for improvement on challenging tasks, such as antibody-antigen docking. These limitations point to clear directions and our future work includes:

- Comprehensive experimental validation across diverse target classes for *de novo* protein design;
- Expansion to nucleic acid and small molecule design tasks, fully leveraging the model’s multimodal capabilities;
- Implementation of advanced inference-time scaling techniques to enhance model performance.

We look forward to collaborating with the scientific community to advance this technology and explore its potential for accelerating discoveries in structural biology and therapeutic development.

## Notes

https://prot.design/

### Competing Interest Statement

The authors have declared no competing interest.

